# Biohybrid restoration of the hippocampal loop re-establishes the non-seizing state in an in vitro model of limbic seizures

**DOI:** 10.1101/2023.01.26.525627

**Authors:** Davide Caron, Stefano Buccelli, Angel Canal-Alonso, Javad Farsani, Giacomo Pruzzo, Bernabé Linares Barranco, Juan Manuel Corchado, Michela Chiappalone, Gabriella Panuccio

## Abstract

**Objective:** The compromise of the hippocampal loop is a hallmark of mesial temporal lobe epilepsy (MTLE), the most frequent epileptic syndrome in the adult population and the most often refractory to medical therapy. Hippocampal sclerosis is found in >50% of drug-refractory MTLE patients and primarily involves the CA1, consequently disrupting the hippocampal output to the entorhinal cortex (EC). Closed-loop deep brain stimulation (DBS) is the latest frontier to improve drug-refractory MTLE; however, current approaches do not restore the functional connectivity of the hippocampal loop, they are designed by trial-and-error and heavily rely on seizure detection or prediction algorithms. The objective of this study is to evaluate the anti-ictogenic efficacy and robustness of an artificial bridge restoring the dialog between hippocampus and EC.

**Approach:** In mouse hippocampus-EC slices treated with 4-aminopyridine and in which the Schaffer Collaterals are severed, we established an artificial bridge between hippocampus and EC wherein interictal discharges originating in the CA3 triggered stimulation of the subiculum so to entrain EC networks. Combining quantification of ictal activity with tools from information theory, we addressed the efficacy of the bridge in controlling ictogenesis and in restoring the functional connectivity of the hippocampal loop.

**Main results:** The bridge significantly decreased or even prevented ictal activity and proved robust to failure; when operating at 100% of its efficiency (i.e., delivering a pulse upon each interictal event), it recovered the functional connectivity of the hippocampal loop to a degree similar to what measured in the intact circuitry. The efficacy and robustness of the bridge stem in mirroring the adaptive properties of the CA3, which acts as biological neuromodulator. Significance. This work is the first stepping stone toward a paradigm shift in the conceptual design of stimulation devices for epilepsy treatment, from function control to functional restoration of the salient brain circuits.

## Introduction

Mesial temporal lobe epilepsy (MTLE) is the most common partial complex epilepsy in adults and the most frequently refractory to medications. While surgical ablation of the epileptic focus is the gold standard to ameliorate drug-refractory MTLE, closed-loop deep brain stimulation (DBS) is the latest frontier treatment [1], holding promise as a less radical approach. Closed-loop DBS relies on seizure detection [2-4] to halt seizures, or, in more advanced paradigms, on seizure prediction [5, 6] to prevent them. The latter is more desirable for ameliorating the patient’s quality of life while minimizing the sequelae of seizures [1]. However, both approaches entail the identification of unique electrographic biomarkers to inform the optimal stimulation timing, and real-time computations that add significant device complexity. Further, the plethora of existing algorithms fail in achieving 100% accuracy and sensitivity [7]. Most importantly, there is still no unifying framework firmly establishing DBS parameters: similar to open-loop DBS, closed-loop devices implement epochs of fixed-frequency stimulation that do not reflect the working mode of the brain and the stimulation policy is established by trial-and-error [8]. As a result, DBS is still unable to guarantee a seizure-free life to each patient: although the median seizure reduction range of 51-70% reported by follow-up studies is remarkable [9], these figures emphasize the inter-individual variability affecting the current clinical scenario and the need of continuing our search for more personalized stimulation strategies. In this regard, the functional restoration of brain circuits compromised by epilepsy would be preferable to the straight modulation of their electrical phenotype pursued by current DBS. To achieve this, the stimulating device should take over the role of the compromised brain pathway(s), thus acting as an integral part of the brain circuit of interest. Bridging neuroprostheses offer this possibility and have already been experimentally demonstrated for sensorimotor recovery after brain damage [10] and spinal cord injury [11]. However, their use for the treatment of epilepsy remains unaddressed. In the particular case of MTLE, the bridging neuroprosthesis should operate the functional restoration of the hippocampal loop. The latter is indeed typically compromised in MTLE, where hippocampal sclerosis is found in >50% of drug-refractory cases, and most often presents with severe damage of the CA1 [12, 13]. This structure accommodates the Schaffer Collaterals, which convey information from the hippocampal subfield CA3 to the entorhinal cortex (EC) via the subiculum located downstream the CA1 and operating as the hippocampal output gate [14]. Thus, the damage of the CA1 disrupts the communication directed from the CA3 to the EC. Noteworthy, in MTLE, the subiculum is typically spared by sclerosis [15-17]. Thus, bridging the CA3 to the subiculum bypassing the damaged CA1 would surrogate the role of the Schaffer Collaterals and re-establish the CA3 output to EC networks via afferent stimulation. Along with surrogating the compromised Schaffer Collaterals, the bridge should convey the appropriate information to restrain ictal activity. In this regard, in vitro studies using rodent hippocampus-EC slices treated with 4-aminopyridine (4AP), where the EC is the primary site of ictogenesis, have pointed to the anti-ictogenic function of the interictal pattern generated by the CA3 [18]. However, this function cannot be exerted if the CA1 is damaged, because the interictal events generated by the CA3 are unable to reach the EC. In this setting, surrogating the frequency of the CA3-driven interictal activity with open-loop periodic stimulation delivered downstream to the Schaffer Collaterals lesion has proved effective in controlling cortical ictal discharges [18, 19]. Building upon these findings, recent work has also demonstrated the efficacy of mimicking, in an open-loop stimulation approach, the temporal dynamics (rather than the average frequency) of the CA3-driven interictal pattern [20], paving the way to an interictal-based grammar for brain stimulation in epilepsy.

In this scenario, a bridging neuroprosthesis relaying the CA3 interictal pattern to the subiculum represents an intriguing and straightforward strategy to restore the functional connectivity of the hippocampal loop and control limbic ictogenesis: it would bypass the need of seizure prediction algorithms while the stimulation policy would be optimized by the real-time feedback from an existing anti-ictogenic interictal pattern rather than by trial and error. Here, we provide the proof-of-concept of this approach using 4AP-treated hippocampus-EC slices in which the Schaffer Collaterals are severed to mimic the CA1 damage often observed in MTLE patients presenting with hippocampal sclerosis. While the bridge operates the artificial reconnection of CA3 to the EC via interictal-driven stimulation of the subiculum, the perforant pathway conveying the biological input of the EC back to the hippocampus is preserved; therefore, this strategy establishes a biohybrid hippocampal loop. We show that this approach effectively restores the CA3-to-EC dialog and controls ictal activity generated by the EC; further, we show that its robustness stems in mirroring the adaptive properties of the CA3. These results demonstrate the possibility for a bridging neuroprosthesis to control limbic seizures that does not depend on seizure detection/prediction algorithms, but on endogenous interictal patterns enabling its intrinsic adaptation to ongoing brain dynamics. By building upon fundamental knowledge of limbic network interactions in MTLE informed by animal models [21, 22] and corroborated by clinical studies [23-25], our work provides a stepping stone toward a new clinically relevant approach for closed-loop DBS in MTLE that outclasses the conceptual design of current devices and paves the way to a paradigm shift from their interaction to their integration within the relevant brain circuits.

## Methods

### Brain slice preparation and maintenance

Horizonal hippocampus-EC slices, 400 μm thick, were prepared from male CD1 mice 4-8 weeks old. Animals were decapitated under deep isoflurane anesthesia, their brain was removed within 60 s, and immediately placed into ice-cold (∼2 °C) sucrose-based artificial cerebrospinal fluid (sucrose-ACSF) composed of (mM): Sucrose 208, KCl 2, KH2PO4 1.25, MgCl2 5, MgSO4, CaCl2 0.5, D-glucose 10, NaHCO3 26, L-Ascorbic Acid 1, Pyruvic Acid 3. The brain was let chill for ∼2 min before slicing in ice-cold sucrose-ACSF using a vibratome (Leica VT1000S, Leica, Germany). Brain slices were immediately transferred to a submerged holding chamber containing room-temperature holding ACSF composed of (mM): NaCl 115, KCl 2, KH_2_PO_4_, 1.25, MgSO_4_ 1.3, CaCl_2_ 2, D-glucose 25, NaHCO_3_ 26, L-Ascorbic Acid 1. After at least 60 minutes recovery, individual slices were transferred to a submerged incubating chamber containing warm (∼32 °C) holding ACSF for 20-30 minutes (pre-warming) and subsequently incubated in warm ACSF containing the K+ channel blocker 4-aminopyridine (4AP, 250 μM), in which MgSO_4_ concentration was lowered to 1 mM (4AP-ACSF, *cf.* [26]). Brain slice treatment with 4AP is known to enhance both excitatory and inhibitory neurotransmission and induce the acute generation of epileptiform discharges [27]. All brain slices were incubated in 4AP-ACSF for at least 60 minutes before beginning any recording session. All solutions were constantly equilibrated at pH = ∼7.35 with 95% O_2_ / 5% CO_2_ gas mixture (carbogen) and had an osmolality of 300-305 mOsm/Kg. Chemicals were acquired from Sigma-Aldrich.

All procedures have been approved by the Institutional Animal Welfare Body and by the Italian Ministry of Health (authorizations 860/2015-PR and 176AA.NTN9), in accordance with the National Legislation (D.Lgs. 26/2014) and the European Directive 2010/63/EU. All efforts were made to minimize the number of animals used and their suffering.

### Microelectrode array recording

Brain slices used for closed-loop stimulation experiments were partially disconnected (*cf*. [28]); namely, the CA3 output to the EC was prevented by Schaffer Collateral’s disruption (either during the slicing procedure or via knife cut), whereas the subicular projections to the EC as well as the EC input to the hippocampus were preserved. Extracellular field potentials were acquired through a 6 × 10 planar MEA (Ti-iR electrodes, diameter 30 μm, inter-electrode distance 500 μm, impedance < 100 kΩ), in which a custom-made low-volume (∼500 μl) recording chamber replaced the default MEA ring (*cf*. [26]) to attain a laminar flow. Individual slices were quickly transferred onto the recording chamber, where they were held down by a custom-made stainless steel/nylon mesh anchor. Slices were continuously perfused at ∼1 ml/min with 4AP-ACSF, equilibrated with carbogen. Recordings were performed at 32° C, achieved with the use of a heating canula (PH01) inserted at the recording chamber inlet port (temperature set at 37° C) along with mild warming of the MEA amplifier base (temperature set at 32° C), both connected to a TC02 thermostat. The recording bath temperature was checked using a k-type thermocouple.

Signals were acquired with either of: (i) the MEA1060 amplifier using the Mc_Rack software, sampled at 2-10 kHz and low-passed at half the sampling frequency before digitization; (ii) the MEA2100-mini-HS60 amplifier using the Multichannel Experimenter software; the amplifier was connected to the IFB v3.0 multiboot interface board through the SCU signal collector unit; signals were sampled at 5 kHz and low-pass filtered at 2 kHz before digitization. In all experiments, signals were stored in the computer hard drive for offline analysis. The custom recording chamber was designed and made by Crisel Instruments, Italy. The equipment for MEA recording was purchased from Multichannel Systems (MCS), Germany.

### Closed-loop stimulation

In all experiments, stimulation was triggered upon detection of the CA3-driven interictal events; this was based on hard-threshold crossing of the signal amplitude, set manually for each experiment at as closely as possible to the signal baseline and no higher than one half of the maximal signal amplitude. Stimuli were delivered in the subiculum and consisted of square biphasic direct current pulses (±100-350 μA, balanced charge, 100 μs/phase, positive phase first) conveyed via MEA electrode pairs connected in bipolar configuration. As the closed-loop system operated in real-time, the delay between interictal event detection and pulse delivery was equal to the sampling interval.

A fast input/output (I/O) curve was always performed prior to the first stimulation protocol to find the current amplitude that would reliably elicit an interictal-like discharge (≥80% response rate, *cf*. [6, 18, 26]). The best performing stimulus amplitude was then kept constant throughout the experiment. Each stimulation session lasted at least 3 times the mean interval of occurrence of ictal discharges (as observed from the preceding control phase) and no less than 20 minutes. Baseline activity in the absence of stimulation was always recorded before and after each stimulation protocol and is overall referred to as control (CTRL).

For the experiments using the MEA1060 amplifier, the closed-loop architecture was designed in Simulink environment (Mathworks, Natick, USA) receiving the MEA signals through a PCI-6255 DAQ board (National Instruments, USA), and interfaced with the MEA1060 system by a custom PCB (PCB Project, Segromigno Monte, Lucca, Italy); in this case, pulses were pre-programmed in the MC_Stimulus II software and downloaded to the STG2004 stimulus generator (both from MCS), which was triggered via TTL. **Supplementary Figure 1** illustrates the Simulink model. The Analog input block reads the MEA signals through the PCI-6255 DAQ board and feeds the CA3 signal from one selected electrode to the Event detection block. The model also implements an Enabling condition for triggering the Stimulation block, to avoid multiple detections of the same event in case the CA3-driven interictal events consisted of brief population busts, and to blank the stimulus artifact that would otherwise lead to a false event detection. Specifically, we set a minimum time window of 250 ms from the last delivered pulse as the enabling condition, based on prior observation of the minimum interval of occurrence and duration of CA3-driven events and the maximum time-lag and duration of the stimulus artifact. Once triggered, the Stimulation block output, in turn, triggers the Digital Output block, which sends a TTL signal to the custom PCB controlling the external stimulator (STG2004, MCS). The Scope visualizes the MEA signals and the output of the Event detection, Time since last event, and Stimulation blocks. For the experiments using the MEA2100-mini, the closed-loop architecture relied on the built-in digital signal processor (DSP) and stimulus generator of the MEA2100 system. The DSP operates with an internal clock of 800 MHZ (1.25 ns period), while the DAC of the stimulus generator operated at 5 kHz (200 μs period).

In a subset of experiments, we challenged the bridge against failure in pulse delivery to mimic missed detections of interictal events (hardware failure) or the CA3 functional impairment described in epileptic tissue [28]. To this end, we set up two different scenarios. In the first scenario, we set a failure rate of 50% (the closed-loop architecture would trigger a pulse every second detected event). To attain this behavior, we have first modified the Simulink model to include a hardware failure module (experiments acquired with the MEA1060 system). As illustrated in **Supplementary Figure 2**, this module is interposed between the Stimulation and the Digital Output block and implements an event counter, which is initialized at 0 and counts up to 1 to attain a 50% failure rate in triggering the Digital Output block. The module also monitors the number of delivered pulses and the number of detected events. Subsequently, this behavior was programmed directly in the built-in DSP of the MEA2100 system. In the second scenario, we set a blanking window of 5 s during which the event detection would not trigger the stimulation. This was implemented in the MEA2100 system as a 0-current tail in the preprogrammed stimulus waveform.

### Data analysis

#### Quantification of epileptiform events

Analysis was performed offline using custom software written in MATLAB R2016b or R2022a. Ictal events were defined based on tonic-like or tonic-clonic-like signal features, disregarding duration cut-off criteria commonly used by other works in the field. Ictal activity was labeled either manually or by means of an automated seizure detection algorithm based on Daubechies discrete wavelet transform type 4 [29], followed by multiple resolution analysis (as in [20]). Detected events were confirmed/corrected by visual inspection as applicable. To take into account the possibility of occurrence of a sole ictal discharge during electrical stimulation, ictal activity was not quantified by its duration and interval of occurrence; rather, we used the ictal state time-percentage (*P*_*ictal*_) as indicator of the overall time spent in the ictal state during each recording phase (*cf*. [6, 30]). *P*_*ictal*_ was computed as in

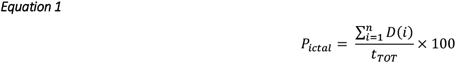

where *i* is ith ictal discharge, *D* is the ictal discharge duration and *t*_*TOT*_ is the total observation time. For CTRL phases (no stimulation), *t*_*TOT*_ is the time between the onset of the first and the termination of the last measured ictal event; for stimulation phases (STIM), *t*_*TOT*_ is the duration of the stimulation protocol. Further, for electrical stimulation, we adopted a conservative approach by including spontaneous or evoked population bursts exhibiting borderline electrographic features that could be classified neither as ictal nor as interictal events; this was done to minimize false negative bias that could yield an overestimation of the stimulation efficacy.

Interictal activity generated by the CA3 was analyzed in terms of statistical parameters **μ, σ** and coefficient of variation (CV = σ/μ) of inter-event interval (IEI) distributions as follows: (i) events timing was retrieved using a threshold-crossing peak-detection; (ii), the interictal IEIs were then fit with a lognormal using the maximum likelihood estimation method, to yield the μ, σ and CV parameters. Lognormal fit was chosen based on the previous demonstration of the lognormal temporal profile of CA3 electrical patterns [20, 31, 32]. To display the distributions, binning used the Freedman-Diaconis algorithm to account for heavy tails. This method was also applied to the analysis of inter-pulse intervals. Dissection of interictal events/pulses belonging to the interictal or ictal periods relied on previously assigned labels of ictal events.

#### Causality analysis

To address the causal relationship between CA3 and EC in the two directions of the loop, we deployed Granger causality (GC), an established approach to quantify how well one time series can be predicted from another[46]. GC explores the system as a linear regression of a stochastic process to measure the amount of information transferred between two signals and yields unique characteristics of signal B inside signal A. To compute the GC, first, we used the Dickey-Fuller test to verify the non-stationarity of MEA signals, which may affect Granger causality; as the signals were indeed non-stationary, they were windowed using a 1-s hamming window to ensure accuracy and avoid artifacts. Then, Vector Auto Regressions (VAR) were fitted to each data window. To establish the model’s best order, we iterated across each window using the Akaike Information Criterion. To ensure the GC calculation will not fail the VAR’s model, residual covariance matrix was checked to be a positive definite. Then, we calculated the GC along the VAR models of each electrode and each window, merging the final GC value in a complete time vector of each electrode, as per **Equation 2**.

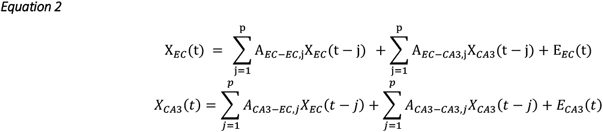

In Equation 2, *p* is the model order, *A* is the matrix containing the contribution of each lagged observation to the predicted values and *E* is the prediction error in each time series (residuals). It can be said that a signal from CA3 “granger causes” a signal from EC if the variance of *E*_*EC*_ is reduced by the inclusion of *X*_*CA3*_; likewise, it can be said that a signal from EC “granger causes” a signal from CA3 if the variance of *EC*_*A3*_ is reduced by the inclusion of *X*_*EC*_.

Analysis was made using custom software written in MATLAB R2022a. Signals were not filtered to avoid artifacts. To compare GC values across different protocols, these were normalized to a range of 0 to 1.

#### Statistical analysis

To compare the efficacy of different stimulation protocols as well as the GC values across multiple groups, we used one-way ANOVA. To this end, we first checked the data for normality (Shapiro-Wilk test) and homoskedasticity (Levene’s test), and we performed the Welch robust test of equality of means (protected ANOVA). Then, we performed pairwise comparisons by either the Games-Howell post-hoc test (unequal variance, equal/unequal sample size) or the Tucky’s post hoc test (equal variance, equal sample size). For statistical comparison between two groups we used the Welch test. Statistical significance set to p < 0.05. Throughout the text, *n* indicates the number of the specified samples; in the figures, the p values are indicated as follows: * p <= 0.05; ** p <= 0.01; *** p <= 0.001. Statistical analysis was performed using custom software written in MATLAB or SPSS 20 (IBM, Armonk, NY, USA).

##### Figures

Figures were prepared in CorelDraw X8. Electrophysiology signals and graphs originally plotted in MATLAB or Microsoft Excel were imported and edited as vector graphics. Violin plots [33] were obtained using the openly shared MATLAB function *violinplot*.*m* (Bechtold, Bastian, 2016. DOI: 10.5281/zenodo.4559847); to preserve the ‘patch’ vector graphics within the violin plots, the MATLAB figures were first converted to pdf via *export_fig*.*m* by Woodford & Altman (https://it.mathworks.com/matlabcentral/fileexchange/23629-export_fig), and then imported in CorelDraw.

## Results

### Concept design of the bridge and establishment of a biohybrid hippocampal loop

Figure 1 illustrates the background knowledge informing the concept design of the bridge (from [18]), and the establishment of a biohybrid hippocampal loop. Each scenario refers to hippocampus-EC slices obtained from healthy rodents and treated with 4AP. When the hippocampal loop is intact, the EC initially generates ictal discharges (not shown, but see [18]); however, these disappear over time due to the anti-ictogenic function of the CA3-driven interictal activity reaching the EC via the Schaffer Collaterals. The schematic rendition depicts the CA3 and EC activities at steady-state, when only interictal discharges are present in both structures. When the hippocampal loop is disrupted by Schaffer Collaterals lesion to mimic the CA1 damage observed in MTLE patients [12], the CA3 output to the EC is hindered, thus permitting the generation of ictal activity (solid line). In this brain slice preparation, the perforant pathway conveying the EC inputs to the dentate gyrus (DG) is still intact; hence, ictal discharges generated by the EC re-enter the hippocampus proper (arrowheads). In the biohybrid loop, the biological perforant pathway co-exists with the artificial reconnection of CA3 and EC operated by the bridge via interictal-driven stimulation of the subiculum (SUB).

**Figure 1.**
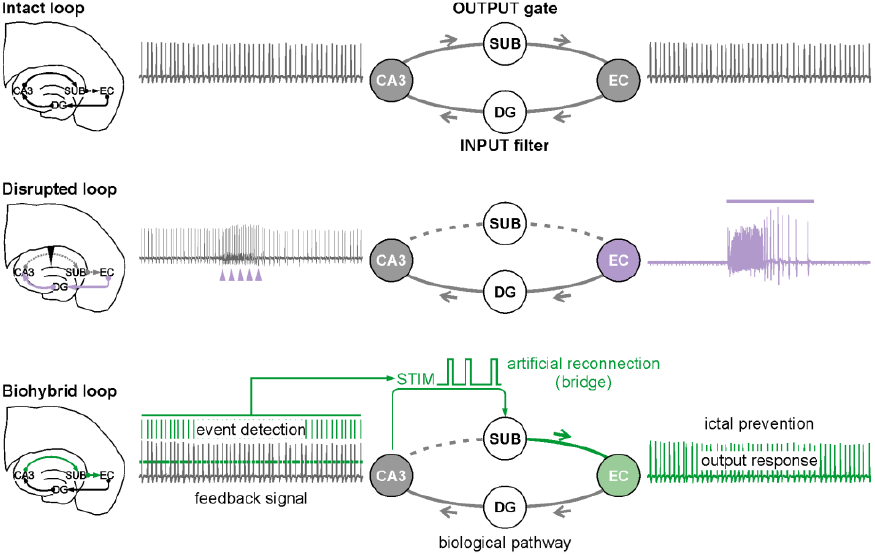
Concept design of the bridge and establishment of a biohybrid hippocampal loop. Schematic rendition of the CA3 and EC interactions during 4AP treatment in the intact, disrupted and biohybrid hippocampal loop brain slices from healthy rodents. On the left is the brain slice schematic representative of the different scenarios. On the right is the corresponding circuit diagram along with the electrical patterns generated by the CA3 and the EC. In the intact hippocampal loop, the CA3 generates a fast interictal pattern that propagates to the EC, controlling its ictogenicity. The representative signals show the electrical pattern recorded at steady-state, i.e., when the CA3-driven interictal activity has eventually abolished ictal discharges originating in the EC. When the hippocampal loop is disrupted by Schaffer Collaterals damage (downward arrow, dashed grey line), the CA3-driven interictal events cannot reach the EC; the latter generates ictal activity (solid line), which invades the hippocampus proper (arrowheads). The core concept design of the biohybrid approach stems in forwarding the CA3-driven interictal pattern to the subiculum via electrical stimulation to recruit and entrain the EC, bypassing the damaged Schaffer Collaterals. As this strategy surrogates the missing CA3-driven signal, it is expected to exert an anti-ictogenic function, thus restoring the scenario depicted for the intact hippocampal loop.

CA3: Cornu Ammonis 3; DG: dentate gyrus; SUB: subiculum; EC: Entorhinal Cortex.

To enable real-time feedback operation of the bridge, early detection of interictal events is here achieved with a simple amplitude threshold-crossing approach, and triggers the stimulation within one sampling interval of threshold crossing. Figure 2 illustrates the real-time operation of the bridge. On the left is a hippocampus-EC slice coupled to a MEA, indicating the feedback, recording and stimulating electrodes. On the right, the signal recorded from the CA3 illustrates the onset of the electrical pulse (STIM, green arrow) upon threshold crossing (green dashed line); the signals from the two indicated cortical locations (CTX1 and CTX2) show the responses evoked by the interictal-driven stimulation of the subiculum.

**Figure 2.**
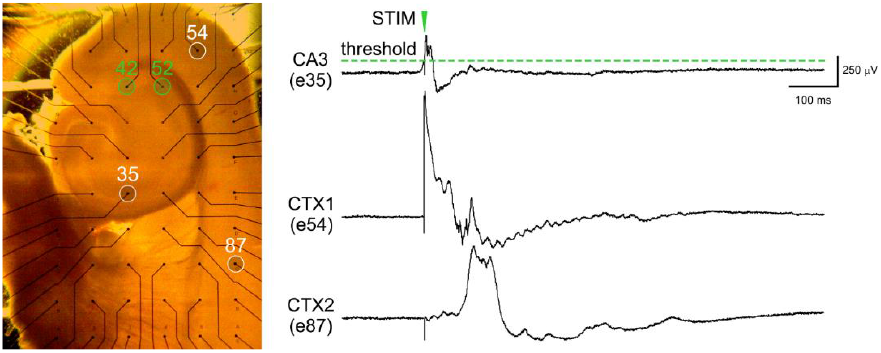
Operation of the bridge. On the left is the picture of a hippocampus-EC slice coupled to a planar MEA, where the white and green circles indicate the recording and stimulating electrodes, respectively. On the right are representative events from the CA3 (feedback signal) and two parahippocampal cortical locations (evoked responses). The threshold value for event detection is set at approximately one half of the signal amplitude (dashed green line). Upon threshold crossing, a pulse is delivered to the subiculum through the stimulating electrode pair, in turn evoking responses in the parahippocampal cortices. Stimulus artifacts are clipped for clarity, and the pulse time is indicated by the green downward arrow.

The reliable operation of the bridge also requires a high signal-to-noise ratio and that interictal events outstand the amplitude of ictal discharges that might occur within the hippocampus proper. Figure 3 illustrates an example of available feedback electrode options (A) and the respective recorded signals (B). In this example, the optimal feedback electrode is e84, as it provides a good signal-to-noise ratio and the interictal events are wider in amplitude than the re-entrant ictal discharges.

**Figure 3.**
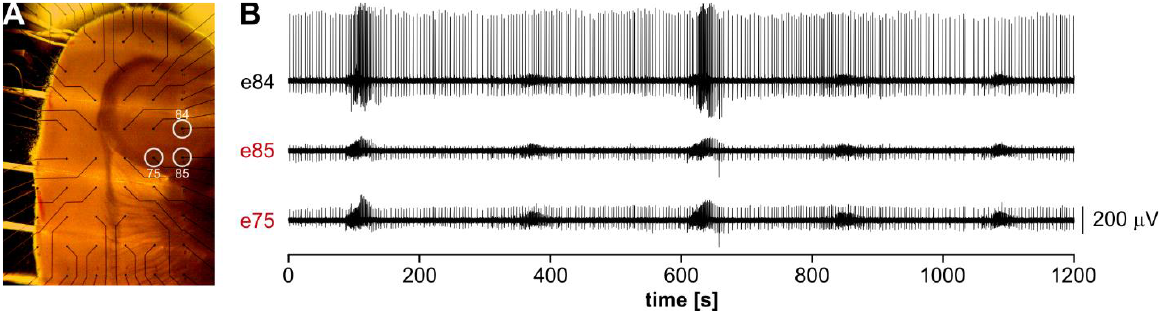
Choice of the optimal feedback electrode. **A**. Picture of a hippocampus-EC slice placed on a 6×10 MEA, showing the set of choices for the CA3 feedback electrode. **B**. Recordings from the electrodes indicated in (A) illustrating optimal (black electrode) and sub-optimal (red electrodes) signal features. In the latter, the signal-to-noise ratio is low and the amplitude of ictal discharges masks the interictal events. The optimal feedback electrode in this representative case is e84 because of the high signal-to-noise ratio and the very small amplitude of ictal discharges compared to the interictal events.

### The bridge is feasible and effective

Figure 4A shows a representative experiment in which the bridge fully prevented ictal activity. This was mostly replaced by brief evoked responses mixed with sporadic population bursts (solid lines). Ictal discharges emerged again upon stimulus withdrawal, confirming that their disappearance was due to the action of the bridge. Figure 4B shows the population bursts pointed by the arrowhead in panel A, visualized at a faster time scale. These events were evoked by the stimulation and exhibited borderline electrographic features that could be classified neither as ictal nor as interictal activity; likely, they reflected temporary hyperexcitability, which, however, could not converge toward a robust ictal discharge because of the ongoing stimulation. As summarized in **Figure 4C**, the bridge proved highly effective, achieving an overall ictal state reduction of 91.31 ± 4.33% (ictal state % – CTRL1: 20.08 ± 1.78, bridge: 1.72 ± 0.81, CTRL2: 20.07 ± 2.19; one-way ANOVA, F(df): 43.98(2), p < 0.001 – bridge vs CTRL1 and CTRL2; n = 9 brain slices).

**Figure 4.**
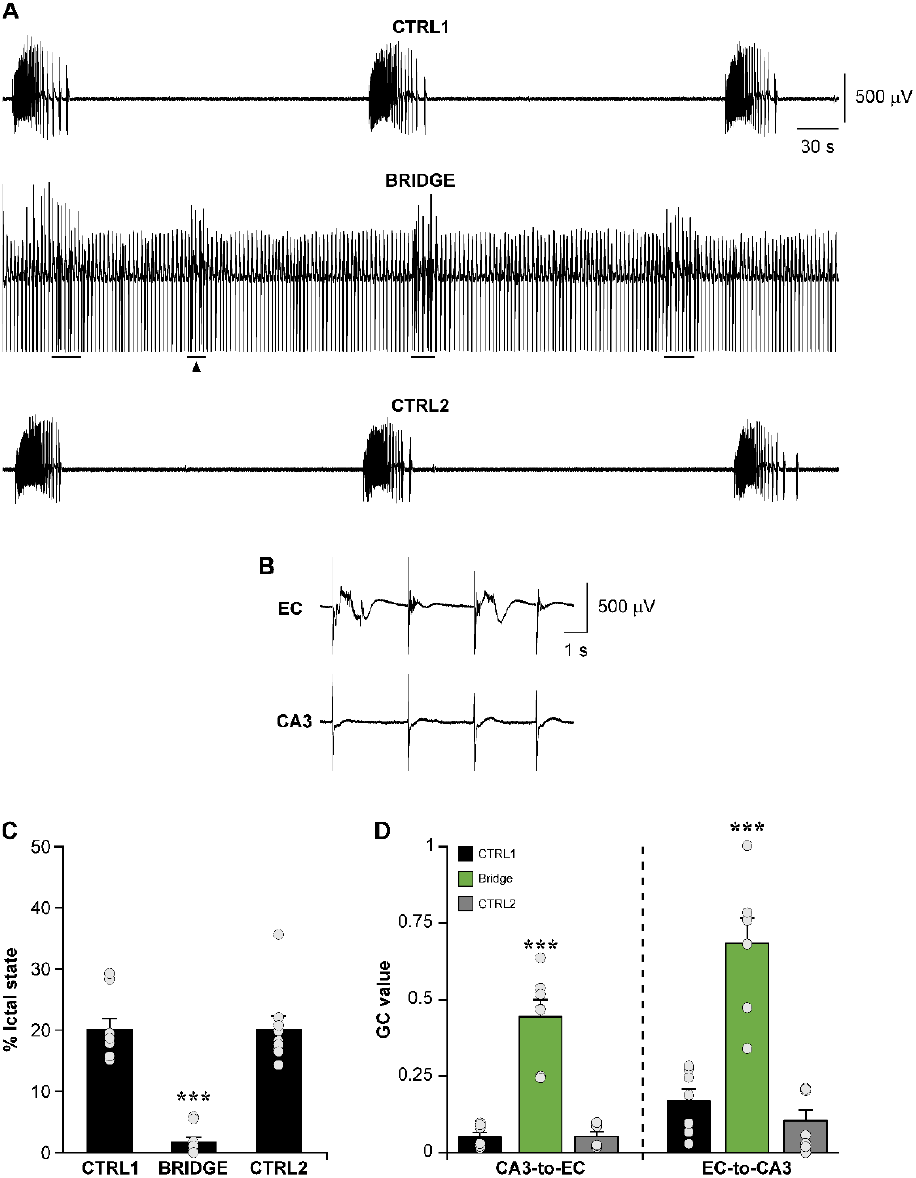
Bridging CA3 and EC controls limbic ictogenesis. **A**. Recordings from the EC during a representative bridging experiment. The pre-stimulus baseline (CTRL1) shows the recurrent generation of ictal discharges; these were abolished by the bridge, and only short epileptiform bursts remained (solid lines). Upon stimulus withdrawal (CTRL2), ictal activity recovered to pre-stimulus condition. **B**. Representative case of epileptiform bursts (pointed by the arrowhead in A) visualized at expanded time scale; the CA3 signal serves as reference for pulse timings. The evoked population bursts likely represented hyperexcitable responses to stimulation. **C**. Summary of the results statistics for this dataset (n = 9 brain slices), showing the high efficacy of the bridge in controlling ictal activity. **D**. Granger causality demonstrate that the bridge induces a strong increase of the functional connectivity between CA3 and EC in the two directions of the loop. In C and D, the dots indicate the values obtained for each brain slice. *** p < 0.001 vs both CTRL1 and CTRL2.

In keeping with the underlying concept of the bridge, we next sought to determine if and to what extent it can restore the functional connectivity of the disrupted hippocampal loop. To this end, we deployed Granger causality (GC) [34], an established method from information theory enabling to address the causal strength between two time series. As shown **Figure 4D**, the GC values demonstrated that the bridge increased the causal relationship between CA3 and EC in the two directions, consistent with the functional restoration of the hippocampal loop (CA3-to-EC – CTRL1: 0.05 ± 0.01; bridge: 0.44 ± 0.06; CTRL2: 0.05 ± 0.02; one-way ANOVA, F(df): 44.47(2), p < 0.001 bridge vs both CTRLs. EC-to-CA3 – CTRL1: 0.17 ± 0.04; bridge: 0.68 ± 0.08; CTRL2: 0.1 ± 0.04; one-way ANOVA, F(df): 31.6(2), p < 0.001 bridge vs both CTRLs).

### The bridge mirrors the adaptive properties of the CA3

The functional restoration of the CA3-to-EC connectivity is the core concept of the bridge. Here, we hypothesized that the CA3 acts as adaptive biological neuromodulator by adapting to the EC drive. To address this hypothesis, we performed GC analysis in hippocampus-EC slices before and after Schaffer Collaterals cut (n = 6 brain slices). **Figure 5A** shows a representative MEA recording from the CA3 and EC of the same brain slice before and after disconnection. The inserts on the right show the boxed signal segments at expanded time scale to emphasize the onset and propagation (or lack thereof) of the interictal and ictal events in the two scenarios.

**Figure 5.**
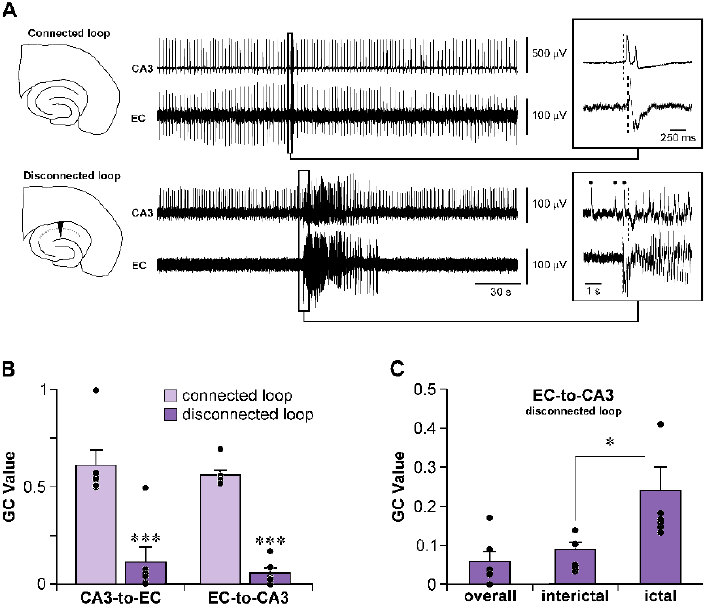
Cortical ictal discharges drive the CA3 in the disrupted hippocampal loop. **A**. Representative epileptiform patterns in the CA3 and EC of a 4AP-treated hippocampus-EC slice before and after disconnection, recorded at steady-state. Connected brain slices generate interictal events only. The insert shows the boxed interictal event at expanded time scale, to emphasize its origin in the CA3 (vertical dashed line). Upon disconnection, the brain slice generates a mixed epileptiform pattern made of interictal and ictal activity. The boxed signal segment frames the ictal onset along with a short pre-ictal period, and is shown in the insert at expanded time scale. Note the disappearance of the CA3-driven events (dots) from the EC and the origin of ictal activity in the latter, with subsequent propagation to the CA3 (vertical dashed lines). **B**. Granger causality analysis of the CA3-to-EC and EC-to-CA3 communication taken as the average GC value across the recorded signal. The significant drop in the GC value upon disconnection is consistent with the disruption of the CA3-to-EC pathway, but not with the preserved EC-to-CA3 connectivity. **C**. Granger causality analysis of the EC-to-CA3 communication in the disconnected loop, dissecting interictal and ictal epochs, evidences the cortical drive during the latter. For clarity, the graph also shows the GC value of the overall signal reported in panel B. In B and C, the dots indicate the values obtained for each brain slice. * p < 0.05; *** p < 0.001.

In connected slices, the GC values confirmed the strong causality between the CA3 and EC, which was similar in the two directions, consistent with the communication loop between these structures (CA3-to-EC: 0.61 ± 0.08; EC-to-CA3: 0.56 ± 0.03; p > 0.05, Welch test). Upon CA3-EC disconnection, the GC values significantly dropped in both directions (CA3-to-EC: 0.11 ± 0.08, p = 0.01 vs connected hippocampal loop; EC-to-CA3: 0.06 ± 0.03, p < 0.001 vs connected hippocampal loop; Welch test; **Figure 5B**).

The drastic reduction of CA3-to-EC causality is expected, since the biological pathway serving it is severed; however, the significant reduction of the EC-to-CA3 causality is surprising, since the perforant pathway is preserved. Possibly, the GC value primarily reflected the interictal periods, since they represented the largest portion of the signal. Thus, we performed a more in-depth analysis of the EC-to-CA3 causality in disconnected brain slices, dissecting interictal and ictal epochs. As shown in Figure 5C, this analysis unveiled the stronger cortical drive during ictal versus interictal periods, with the latter being statistically similar to what measured for the overall signal (GC value – ictal: 0.2 ± 0.05, interictal: 0.07 ± 0.02, overall: 0.06 ± 0.03; ANOVA, F(df): 6.78(2), p < 0.05 ictal vs both interictal and overall signal; p > 0.05 overall signal vs interictal). Next, we analyzed the temporal distributions of the CA3-driven interictal events in an extended dataset of 59 disconnected brain slices, again dissecting ictal from interictal epochs. **Figure 6A** shows a representative recording from the CA3 and EC and the distribution parameters (μ, σ, and CV = σ/μ) of interictal inter-event intervals (IEI) demonstrating the acceleration of the CA3-driven interictal events during the ictal periods. As summarized in **Figure 6B**, this behavior was overall confirmed by population statistics (μ interictal: 1.84 ± 0.14, μ ictal: 1.38 ± 0.1, p < 0.01; σ interictal: 1.30 ± 0.02, σ ictal: 1.4 ± 0.03, p > 0.01; CV interictal: 0.89 ± 0.05, CV ictal: 1.33 ± 0.1, p < 0.001; Welch test).

**Figure 6.**
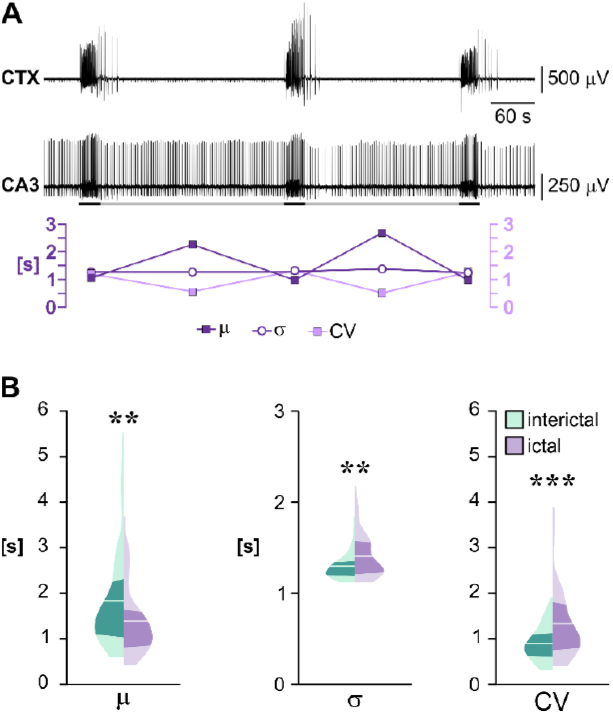
The CA3 is an adaptive biological neuromodulator. **A**. Representative recording from EC and CA3 of a disconnected brain slice showing the acceleration of the CA3-driven interictal events upon entrainment by ictal activity. The black and grey lines below the CA3 signal mark the ictal and interictal epochs, respectively, and serve as reference for the statistical parameters of the CA3 interictal temporal profile plotted below the signal. The decreased μ and CV substantiate the interictal acceleration. **B**. Violin plots summarizing the results statistics for the interictal temporal profile of the analyzed dataset (n = 59 brain slices), demonstrating the consistent acceleration of interictal events during epochs of ictal entrainment. The solid lines indicate the mean; the darker areas frame the quartiles. ** p < 0.01; *** p < 0.001.

These results anticipate that the CA3 may act as adaptive biological neuromodulator: while the disruption of the Schaffer Collaterals impede the CA3 from relaying its drive to the EC, the CA3-driven stimulation will adapt to the evolving network dynamics. To corroborate this point, the ideal experimental scenario would be a bridge that gradually dampens rather than prevents ictal activity. This would enable to analyze the temporal distribution of the delivered pulses (hence of the CA3-driven interictal events) during the progressive abatement of ictal discharges. **Figure 7A** shows such a case: ictal discharges became gradually shorter until they were replaced by evoked interictal-like events (events of interest are highlighted in light grey in the EC signal). As indicated by the pulse times (green bars), the CA3-driven stimulation accelerated during these discharges; further, the distribution parameters of the inter-pulse intervals initially exhibited the oscillatory trend of interictal-to-ictal transitions (*cf*. **Figure 6**); eventually, they stabilized to values similar to the ictal-free periods. The transition from the last population burst to the stable pattern of interictal-like evoked discharges is shown at expanded time scale in **Figure 7B**. Statistical analysis of the temporal profile of the delivered stimuli confirmed the accelerating behavior of the bridge during the highlighted epochs (**Figure 7C**). In our hands, we encountered only one such case, because the bridge invariably exerted a strong anti-ictogenic effect. Nonetheless, this scenario suggests that the efficacy of the bridge stems in its inherent feature of relaying the CA3 adaptation to EC inputs back to the EC.

**Figure 7.**
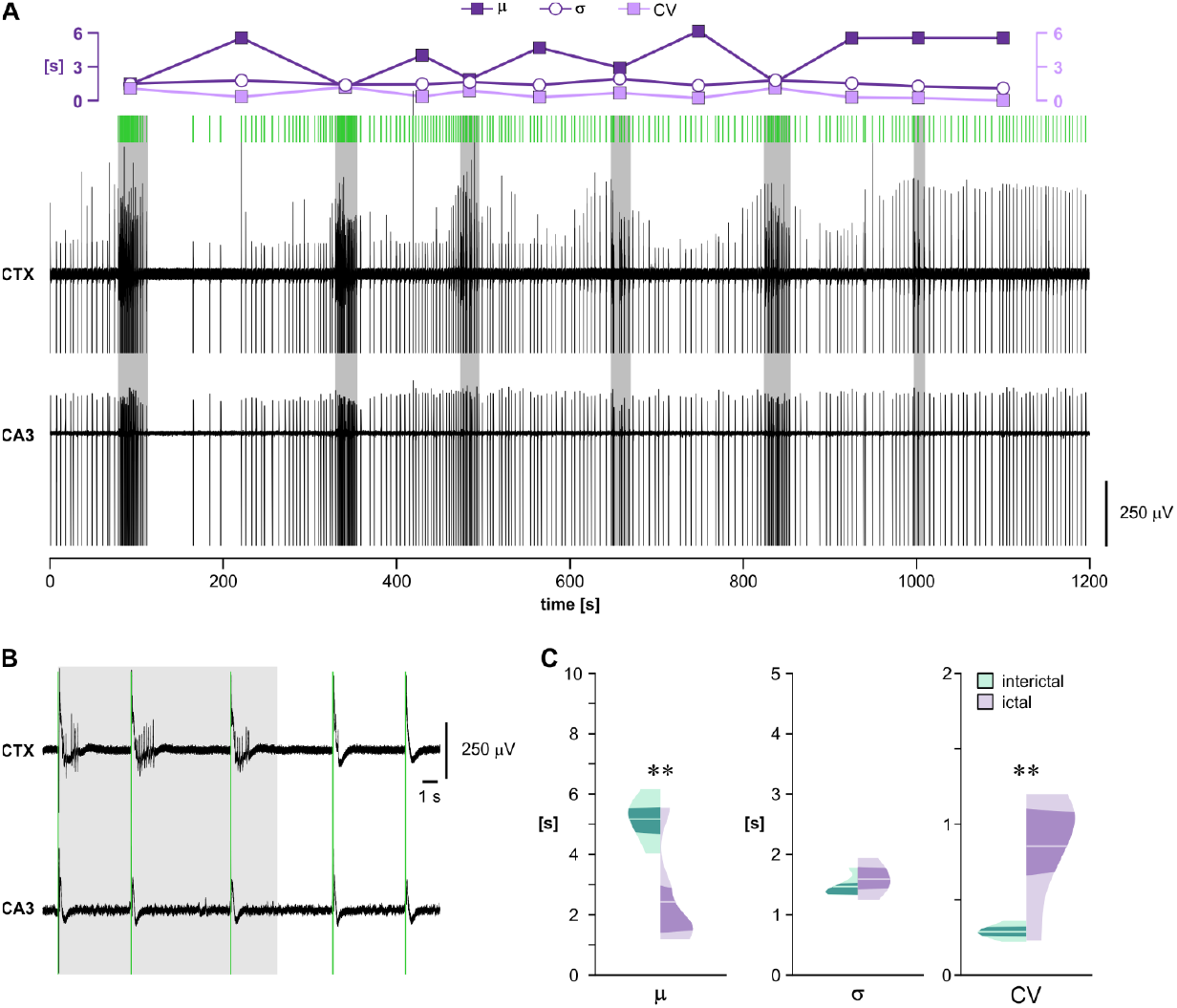
The bridge mirrors the adaptive properties of the CA3. **A**. Full recording of a representative 20-minute experiment in which the bridge gradually abolished ictal activity. Stimulus artifacts are not blanked to demonstrate the operation of the bridge. Epileptiform bursts (highlighted by the grey shading) gradually transitioned from robust ictal discharges to interictal-like evoked responses. The green bars above the signal mark the timings of the delivered pulses to emphasize the emerging accelerations in the CA3-driven stimuli. The insert above the signal shows the computed statistical parameters of the temporal distribution of the pulses, starting from the first labeled event: while the σ parameter remained stable over time, the μ parameter markedly decreased during these events; likewise, the CV parameter was higher. At the time of the last and shortest labeled event, the temporal distribution of the pulses settled to values similar to ictal-free periods. **B**. Recording segment starting from the onset of the last burst discharge in A, visualized at a faster time scale, demonstrating the entrainment of EC networks by the ongoing stimulation and transition of evoked responses from population bursts to interictal-like discharges. Pulse timings are indicated by the green bars. C. Summary of the result statistics for this experiment, confirming the adaptive behavior of the bridge. ** p < 0.01.

### The bridge efficacy relies on the temporal statistics of the CA3-driven interictal pattern

The CA3 is often compromised in MTLE patients with hippocampal sclerosis [12]; in animal models, this relates with a decreased frequency of the CA3-driven interictal discharges, which is approximately halved compared to the non-epileptic control [28], posing the question of the efficacy of the bridge in such cases. Further, DBS hardware malfunction is a common cause of reappearance or worsening of symptoms in the clinical setting (15-30% of the cases [35]). To address the bridge robustness to such conditions, we challenged it against a failure rate of 50% (i.e., the stimulator delivered a pulse every second detected event, **Figure 8A**); this paradigm is hereafter referred to as bridge-50%. In these setting, the fully efficient bridge, hereafter referred to as bridge-100%, served as positive control, and the stimulation protocols were shuffled. **Figure 8B** shows the recordings from the EC in a representative experiment; for simplicity, we only show the bridge-50% and its related CTRL phases. In this experiment, the bridge-50% prevented the generation of ictal discharges. Only a short epoch of stimulus-independent population bursts could be observed early during the stimulation (solid line), which was eventually entrained (**Figure 8C**). As summarized in **Figure 8D**, the bridge-50% performed similarly to the bridge-100% (n = 7 brain slices; ictal state % – pooled CTRLs: 17.67 ± 1.39; bridge-50%: 1.06 ± 0.83; bridge-100%: 0.8 ± 0.69; ictal state reduction vs preceding CTRL – bridge-50%: 92.38 ± 6.51%; bridge-100%: 95.2 ± 4.57; one-way ANOVA, F(df): 43.8(2); p < 0.001 for both bridges against the pooled CTRLs; p > 0.05 between the two bridges). Further, the effective stimulation frequency (n. pulses/stimulation duration) exhibited a variable range (0.13-0.52 Hz), which did not correlate with the degree of ictal reduction (**Figure 8E**); the mean frequency was 0.3 ± 0.06 Hz with frequencies < 0.2 Hz in ∼43% of the cases.

**Figure 8.**
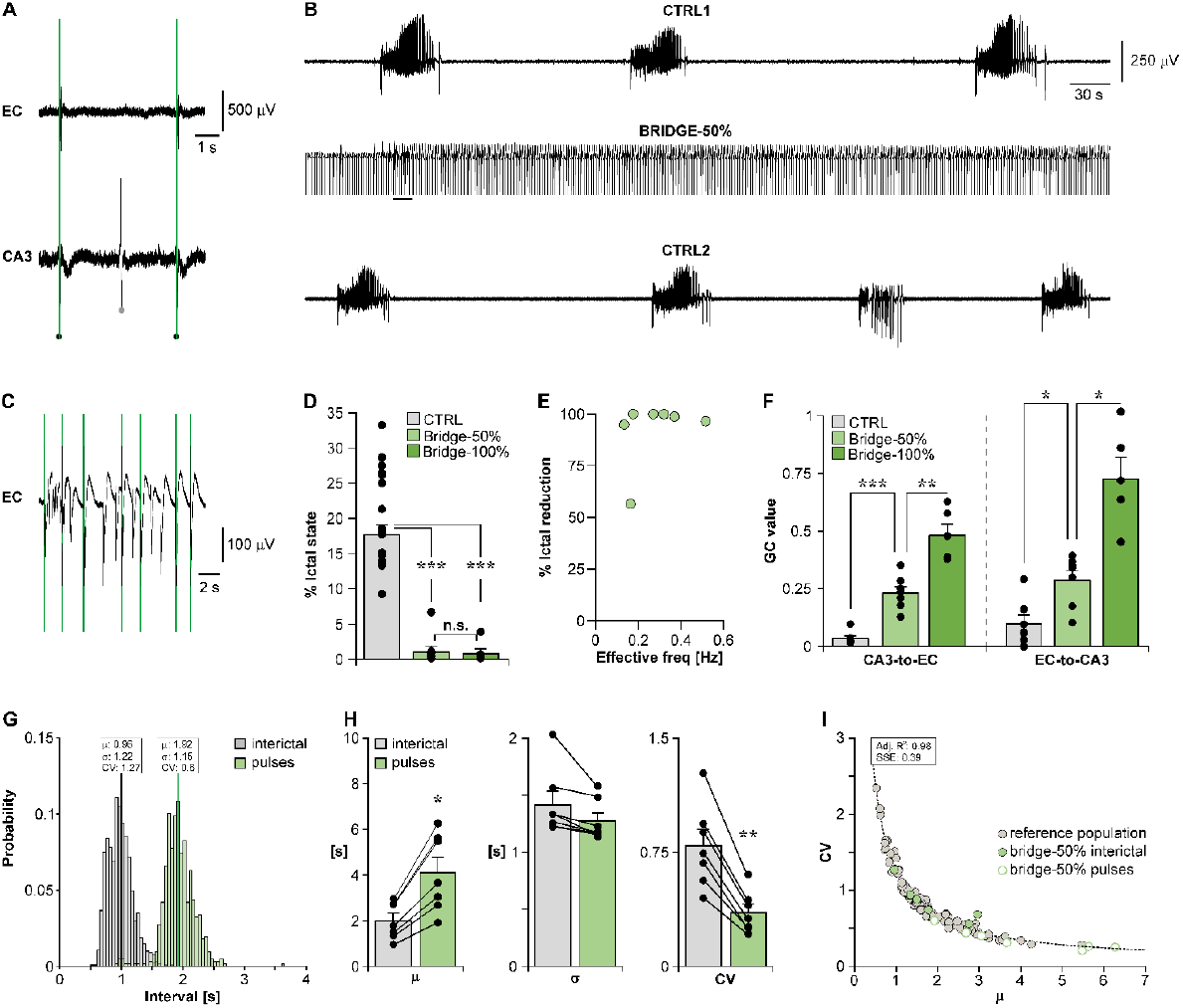
Bridge robustness to failure suggests a pulse-timing and pattern dependent efficacy. **A**. Representative recordings from EC and CA3 illustrating the operating mode of the bridge-50%. The black dots indicate the detected interictal events that triggered the stimulation, while the grey dot indicates the detected interictal event that did not trigger it. The vertical green bars mark the pulse timings. **B**. Recording from the EC during a representative experiment. The bridge abolished ictal activity: only one short population burst was observed initially (solid line), but was eventually entrained by the stimulation. **C**. Population burst indicated by the solid line in B, visualized at expanded time scale. Vertical green bars mark the pulse timings. **D**. Summary of the results statistics obtained from this dataset (n = 7 brain slices) demonstrating the similar performance of the bridge-50% and the bridge-100%. **E**. The efficacy of the bridge-50% does not correlate with its effective stimulation frequency (n. pulses/stimulation duration). F. Granger causality reveal the partial restoration of the functional connectivity of the hippocampal loop in both directions, compared to the bridge-100%. **G**. Distributions of the interictal and inter-pulse intervals for the experiment in B show that the inter-pulse distribution is right-shifted compared to the interictal distribution, while the shape is preserved. **H**. Comparison of the distribution parameters μ, σ and CV between interictal and pulse intervals: in the latter, the μ parameter is ∼double and accounts for the ∼halved CV. **I**. The μ vs CV relationship of the inter-pulse intervals of the bridge-50% fits the distribution of the original interictal events and of a larger interictal population of reference (n = 105 brain slices, same dataset as in[20]), indicating that the inter-pulse intervals represent a subset of the same population. In D, F and H, the dots indicate the values obtained for each brain slice. * p < 0.05; ** p < 0.01; *** p < 0.001.

Granger causality **(Figure 8F)** showed that the bridge-50% improved the functional connectivity of the hippocampal loop in both directions; however, consistent with a 50% failure, the achieved functional restoration was approximately half of what achieved by the bridge-100% (GC value CA3-to-EC – CTRL: 0.03 ± 0.01, bridge-50%: 0.23 ± 0.03, bridge-100%: 0.48 ± 0.05; ANOVA F(df): 56.5(2), p < 0.001 vs CTRL; p < 0.01 vs bridge-100%. GC value EC-to-CA3 – CTRL: 0.1 ± 0.04, bridge- 50%: 0.29 ± 0.09, bridge-100%: 0.72 ± 0.09; ANOVA F(df): 30.7(2); p < 0.05 vs CTRL and bridge-100%.). Yet, the ictal reduction similar to the bridge-100% and the frequency-independent efficacy indicates robustness to failure.

To further address this phenomenon, we analyzed the statistical properties of the inter-pulse intervals during the bridge-50%. **Figure 8G** shows the probability distributions for the representative case shown in panel **B**; as expected, the inter-pulse interval distribution is right-shifted and the μ parameter is doubled compared to the distribution of the interictal IEI; remarkably, the distribution shape is fairly preserved. Comparison of the μ, σ, and CV parameters of pulse versus interictal intervals across the dataset (**Figure 8H**) confirmed the two-fold increase in the μ parameter (interictal: 2 ± 0.33, pulses 4.1 ± 0.69, p < 0.05, Welch test) and indicated the statistical similarity of the σ parameter (interictal: 1.41 ± 0.12, pulses 1.27 ± 0.07, p > 0.05, Welch test); in keeping with this, the CV was approximately halved (interictal: 0.79 ± 0.11, pulses: 0.35 ± 0.05, p < 0.01, Welch test).

Previous analysis of a large interictal population has shown that the μ vs CV relationship of their intervals follows a power-law [20]; if the inter-pulse intervals in the bridge-50% represent a sub-population of the interictal intervals, their μ vs CV relationship should follow the same power-law. **Figure 8I** demonstrates that both interictal events and delivered pulses fit within the same power-law relationship of the reference population (from [20]); in fact, their CV was not significantly different from what estimated from the μ vs CV fitting parameters (empirical CV: 0.35 ± 0.05, estimated CV: 0.41 ± 0.07; p > 0.05, Welch test). Thus, the bridge robustness to failure may reside in the preserved temporal statistics of the relayed interictal events.

As a counterproof, we performed additional experiments in which the interictal temporal statistics were dissipated by introducing a 5-s blanking window during which the event detection would not trigger the stimulator; in this setting, the failure rate varies between pulses depending on the number of interictal discharges occurring during the set time constraint. This paradigm is hereafter referred to as bridge-bw5. The duration of the blanking window was chosen based on two observations: (i) the bridge-50% outperformed the limit open-loop stimulation frequency of 0.2 Hz (*cf*. **Figure 8D**); (ii) the μ vs CV power-law relationship of interictal events tends to its asymptote at μ ≥ 5 s, i.e., at frequencies ≤ 0.2 Hz (*cf*. **Figure 8I**); herein, the CV reaches its lowest values tending toward a periodic pattern.

**Figure 9A** shows the variable interictal-to-pulse failure rate in a representative case. Overall, the failure rate ranged between 72.73 and 84.64% (79.32 ± 1.32%; n = 6 brain slices). In this series of experiments, again, the bridge-100% was used as positive control and the stimulation protocols were shuffled.

**Figure 9.**
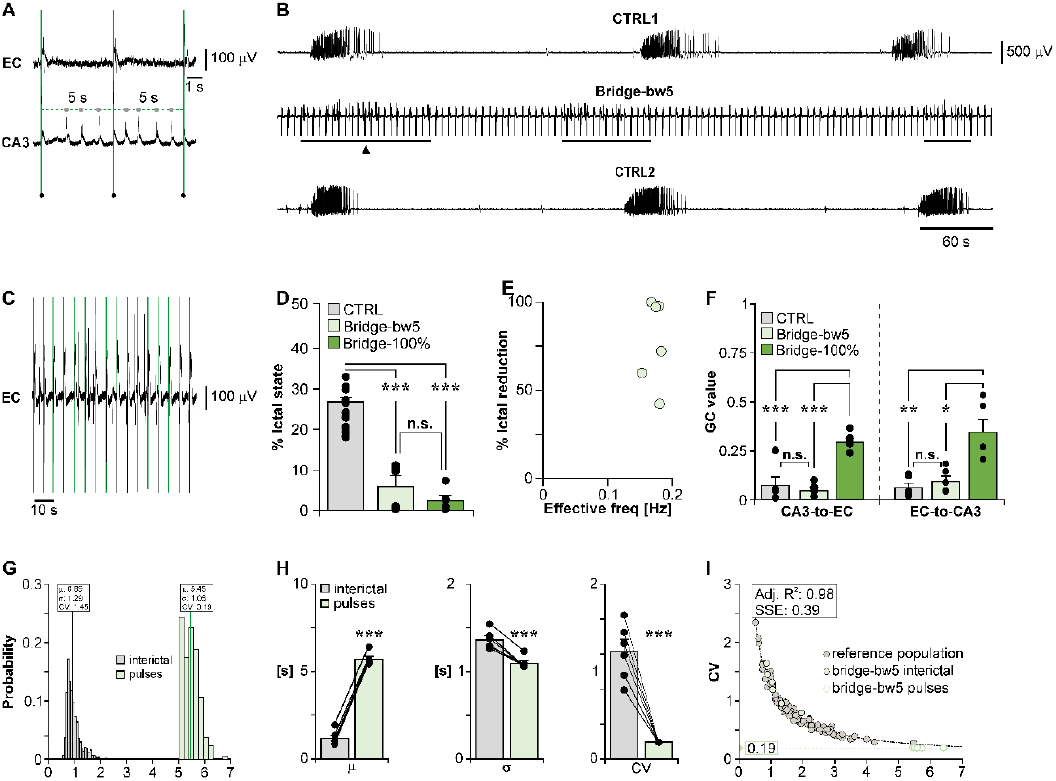
The bridge performs unreliably upon dissipation of the interictal temporal statistics in the pulse timing. **A**. Representative recordings from the EC and CA3 illustrating the operating mode of the bridge-bw5. The black dots indicate the detected interictal events that triggered the stimulation, while the grey dots indicate the detected events that fell within the 5-s blanking window. The vertical green bars mark the pulse timings. **B**. Recording from the EC during a representative best-case experiment: the bridge abolished ictal activity and the only epileptiform events were stimulation-independent interictal-like discharges (solid lines). **C**. Representative case of stimulus-independent activity corresponding to the epoch pointed by the arrowhead in B, visualized at expanded time scale. The vertical green bars mark the pulse timings. **D**. Summary of the results statistics obtained from this dataset (n = 6 brain slices) showing a statistically similar efficacy of the bridge-bw5 and the bridge-100%. **E**. The efficacy of the bridge-bw5 is highly variable and does not correlate with its effective stimulation frequency (n. pulses/stimulation duration). **F**. Granger causality confirms the poor performance of the bridge-bw5 in re-establishing the functional connectivity of the hippocampal loop, which does not differ from CTRL. **G**. Distributions of the interictal and inter-pulse intervals for the experiment in B: the inter-pulse distribution is right-shifted and its shape is unpreserved compared to the interictal distribution. **H**. Each of the distribution parameters μ, σ and CV are different between interictal and pulse intervals. **I**. The μ vs CV relationship of the interictal IEI falls within the reference distribution (n = 105 brain slices, same dataset as in [**20]**); however, the inter-pulse intervals of the bridge-bw5 fall below the asymptote of the reference fitted curve, indicating that they do not represent a subset of the same population. The CV values approximate a constant of 0.19 (dashed line), which is in contrast with the inter-individual variability observed in interictal patterns. In D, F and H, the dots indicate the values obtained for each brain slice. * p < 0.05; ** p < 0.01; *** p < 0.001.

**Figure 9B** shows a representative best-case scenario in which the bridge-bw5 prevented ictal activity; cortical networks were fully entrained by the stimulation and only generated short epochs of stimulus-independent interictal-like events (solid lines; epoch marked by the arrowhead is expanded in **Figure 9C**). As summarized in **Figure 9D**, the bridge-bw5 overall performed similarly to the bridge-100% (ictal state % – pooled CTRLs: 23.61 ± 1.04, bridge-bw5: 5.25 ± 2.36, bridge-100%: 2.1 ± 1.19; ictal state reduction vs preceding CTRL – bridge-bw5: 77.59 ± 10.73%; bridge-100%: 90.33 ± 6.32; one-way ANOVA, F(df): 79.9(2), p < 0.001 for both bridges against the pooled CTRLs; p > 0.05 between the two bridges). However, the bridge-bw5 performance was highly variable, ranging 44.63-100% of ictal reduction, and achieving full efficacy in 50% of the cases only (**Figure 9E**). Further, Granger causality evidenced no significant change in the CA3-to-EC correlation in both directions of the loop (**Figure 9F**; GC value CA3-to-EC – CTRL: 0.07 ± 0.04, bridge-bw5: 0.05 ± 0.01, bridge-100%: 0.29 ± 0.02; ANOVA F(df): 34.1(2), bridge-bw5 vs CTRL: p > 0.05, bridge-bw5 vs bridge-100%: p < 0.001, bridge-100% vs CTRL: p < 0.001. GC value EC-to-CA3 – CTRL: 0.06 ± 0.02, bridge-bw5: 0.09 ± 0.03, bridge-100%: 0.35 ± 0.07; ANOVA F(df): 16.5(2); bridge-bw5 vs CTRL: p > 0.05, bridge-bw5 vs bridge-100%: p < 0.05, bridge-100% vs CTRL: p < 0.01).

The variability in ictal state reduction and lack of causality restoration is consistent with the misalignment of the bridge with the temporal evolution of the CA3-EC dynamic interplay as a consequence of the constrained inter-pulse interval. In keeping with this, not only the distribution of the inter-pulse intervals was right shifted compared to the original interictal IEI distribution; its shape did not seem preserved (**Figure 9G**; data from representative case in panel B). Comparison of the μ, σ, and CV parameters of pulse versus interictal intervals in the dataset evidenced the significant change in each of them (**Figure 9H**; μ − interictal: 1.19 ± 0.18, pulses 5.7 ± 0.16, p < 0.001; σ − interictal: 1.36 ± 0.05, pulses 1.09 ± 0.03, p < 0.001; CV – interictal: 1.23 ± 0.14, pulses: 0.19 ± 0.001, p < 0.001, Welch test).

Further, as shown in **Figure 9I**, the μ vs CV relationship of the delivered pulses fell below the asymptote of the reference curve and the CV parameter approximated a constant value (0.19 ± 0.001) which differed significantly from what estimated from the fitting parameters of the reference curve (0.27 ± 0.006, p < 0.001, Welch test).

**Figure 10** provides the performance overview for the different bridging paradigms, including ictal reduction percentage **(A)**, and the achieved functional restoration of the hippocampal loop compared to connected/disconnected brain slices, taken as positive and negative control, respectively **(B)**. Remarkably, the GC values show a clear trend toward increased functional connectivity with the increased bridge efficiency (see **Supplementary Table 1** for full results statistics).

**Figure 10.**
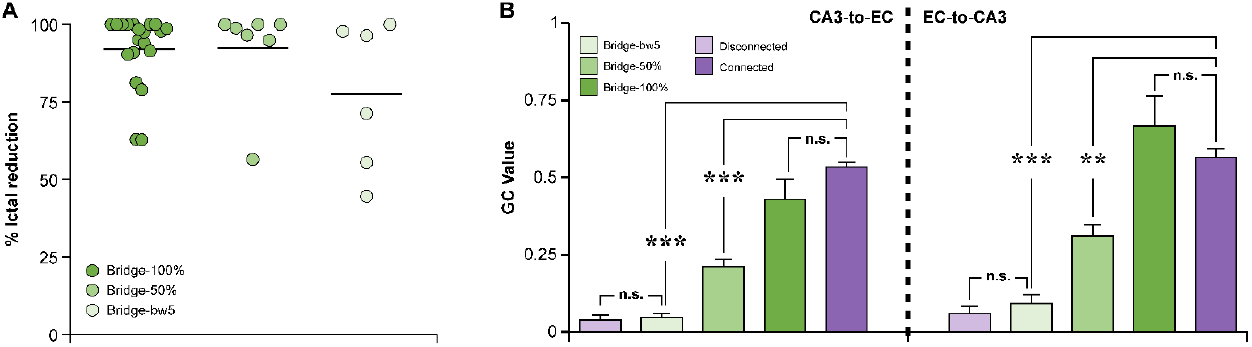
Performance overview of the bridge under different failure testing paradigms. **A**. Direct comparison of ictal state reduction achieved by the three bridging paradigms. Each dot represents one experiment, while the solid line indicates the mean for each dataset. **B**. Granger causality indicates that the degree of functional restoration of the hippocampal loop follows the efficiency of the bridge in forwarding the CA3-driven interictal events to the subiculum. The bridge-100% attains GC values similar to what measured in the intact hippocampal loop; conversely, the bridge-bw5, which dissipates the temporal statistics of the CA3-driven interictal pattern, exhibits GC values comparable to those of the disconnected hippocampal loop.

## Discussion

Closed-loop DBS is the latest frontier in neuromodulation to ameliorate drug-refractory epilepsy, as it provides ad hoc intervention based on the patient’s brain activity. However, it still relies on arbitrary settings defined by trial-and-error, and it is primarily designed to halt rather than prevent seizures. In addition, delayed stimulation is a drawback of current closed-loop DBS, because this is based on detection of abnormal brain activity within a sliding window, using complex signal processing methods [36, 37]. As a result, the median seizure reduction rate reported by follow-up evaluations is a remarkable yet improvable 51-70% [9] element. This highlights the need of pinpointing the most appropriate stimulation timing and of rooting pulse sequences in physiologically meaningful patterns, providing timely personalized interventions to prevent rather than halt seizures. In turn, this requires a paradigm shift in stimulation therapies for epilepsy, namely shifting their conceptualization from interaction with to integration within the brain. In fact, current DBS devices are conceived to rectify pathological brain activity via unnatural interventions, rather than to replace a compromised brain function or pathway via a brain-inspired operating mode. The latter requires the circuit-informed device design that is offered today by neuroprosthetics acting as a bridge between disconnected brain areas. This approach goes beyond the canonical acceptance of DBS as external intervention. Remarkably, bridging neuroprostheses have already shown promising results in pre-clinical studies aimed at the recovery of sensorimotor function after brain damage [10] and spinal cord injury [11]. The same principle can thus be applied to epileptic syndromes when the underlying circuitry is known and well characterized. In this regard, animal models complement human studies by providing precious information that could not be acquired otherwise. In the specific case of MTLE, the large body of evidence obtained with the help of animal models has enabled a more in-depth understanding of the network interactions underlying it [22]. In this work, we have built upon this knowledge and we have used an established in vitro model of limbic ictogenesis to demonstrate the proof of principle of an artificial bridge to restore the functional connectivity of the hippocampal loop and so restrain limbic ictal activity. This model offers several parallelisms with human MTLE: **(i)** the brain slice preparation includes the fundamental limbic circuits involved in this epileptic syndrome; **(ii)** treatment of this brain slice preparation with 4AP induces the acute generation of interictal and ictal epileptiform discharges exhibiting electrographic features similar to what seen in human MTLE; **(iii)** the Schaffer Collaterals disruption mimics the damage of the CA1 hippocampal subfield often observed in drug-refractory MTLE patients presenting with hippocampal sclerosis; **(iv)** the primary origin of ictal activity in the EC reflects clinical findings reporting the frequent seizure onset in this cortical region [22-25] and the benefit of removing it alone or in combination with the hippocampus to treat drug-refractory MTLE [22].

The results of the present study demonstrate the feasibility and efficacy of the CA3-to-EC bridge not only in controlling ictal activity, but also in restoring the functional connectivity of the hippocampal loop to a similar extent of the intact one. The bridge controlled ictal activity similarly well to what reported by open loop stimulation studies deploying periodic [18, 20] or distributed pulse trains [20] reflecting, respectively, the mean frequency or temporal distribution of the CA3-driven interictal events. Remarkably, it also performed well when the effective stimulation frequency was below or equal to 0.2 Hz, a periodic stimulation frequency that was previously shown to exhibit sub-optimal and unreliable seizure suppression both in vitro [19, 30, 38] and in vivo [39], regardless of the stimulation target. The results of the present study also support the view that the efficacy and robustness of the bridge stem in mirroring the adaptive properties of the CA3. This hippocampal region has been widely studied for its pacemaker-like features [40, 41] and its anti-ictogenic function in limbic circuits involved in MTLE [18, 42-44]. Our work provides the first evidence that it acts as adaptive biological modulator of cortical hyperexcitability (**Figure 6**). Namely, we have shown that cortical ictal activity invading the hippocampus proper drives the CA3, which responds with accelerations of interictal events, likely aimed at halting the seizure (**Figure 6**). By relaying the adaptive behavior of the CA3 back to the EC (**Figure 7**), the bridge effectively and reliably controlled ictal activity as long as the intervals of the delivered pulses reflected the temporal evolution of the CA3-EC dynamic interplay. In fact, the bridge demonstrated robust to 50% failure but still abiding to the temporal statistics of the CA3-driven interictal pattern, a paradigm primarily mimicking the CA3 functional impairment possibly accompanying MTLE [28]; at variance, the bridge-bw5, in which the CA3 interictal temporal dynamics where dissipated by the stimulation to primarily mimic hardware malfunction, yielded an unreliable performance.

Granger causality analysis provided further insight into the intrinsic adaptive properties of the bridge and its ability to restore the functional connectivity of the disrupted hippocampal loop (**Figure 10**): the recovery of causality to a statistically similar level to what measured in the intact hippocampal loop along with the restraint of ictal activity suggests that the bridge operates according to the most appropriate stimulation timing. In fact, network responses to perturbations are state-dependent [45, 46]; thus, if the stimulated network is in a refractory state, the stimulus will not evoke a response. Interictal discharges themselves are to be regarded as a perturbation to the system [46]; as such, they may also induce refractoriness [23, 47] and so exert an anti-ictogenic role [46]. We may therefore hypothesize that in establishing a biohybrid hippocampal loop, the biological and artificial counterparts engage in a reciprocal dynamic adaptation, informed by the CA3, and settle to a new steady-state, free of seizure activity (**Figure 7**). In this view, the advantage of the bridge over other stimulation paradigms is two-fold: **(i)** the stimulation policy is provided directly from the epileptic brain, thus surpassing the long-standing issue of trial-and-error; **(ii)** the functional reconnection of disconnected brain areas goes beyond the current concept of DBS as external perturbation of pathological brain dynamics.

When a circuit-informed bridging neuroprosthesis is not feasible, interictal discharges may still present the key for an effective closed-loop DBS, as they reflect the propensity of the epileptic brain to generate seizures [48-50]. Nonetheless, the concept of an interictal-informed closed-loop DBS has only recently emerged [51, 52] and the direct exploitation of interictal discharges to drive the closed-loop stimulation has never been considered prior to the present work. This is because a clear definition of their pro-ictogenic versus anti-ictogenic role is still a matter of debate due to the general assumption that the two roles are mutually exclusive. In contrast, emerging evidence strongly indicates that such dichotomic definition is invalid; rather, it points to the co-existence of these opposite roles as state-dependent features in the bistable dynamical system that is the epileptic brain [46, 48-50]. In this view, and as also discussed in [52], interictal discharges may be exploited to probe ongoing brain dynamics (seizure prediction) and so inform the stimulation timing. Along these lines, the present work demonstrates the proof-of-concept for two distinct yet complementary aspects for biomedical devices for epilepsy treatment: (i) a bridging neuroprosthesis reconnecting brain areas whose impaired functional dialogue leads to seizure generation (here, the CA3 and the EC), and (ii) an interictal-based grammar for closed-loop DBS.

Although the brain slice preparation used in this study is a reductionist model of the limbic circuitry involved in MTLE and only permits acute studies in the absence of a behavioral correlate, the evidence obtained in this simplified in vitro setting is a fundamental stepping stone to inform pre-clinical experiments in epileptic rodents toward the clinical translation of the proposed approach. Therefore, the next step will be evaluating the long-term anti-ictogenic effect of the bridge as well as its effects on cognitive function, which is often impaired in MTLE [53].

The potential translation of the proposed approach to the clinical setting may benefit those patients who do not present with a severe neuronal loss in the CA3. The latter is most common in hippocampal sclerosis (HS) type I (as per the ILAE consensus classification [13, 54]), although the reported degree of neuronal loss ranges from mild (HS type Ia) to severe (HS type Ib) across different cases. Remarkably, the CA3 is relatively spared in other HS types. However, it needs to be noted that the exact incidence of hippocampal sclerosis (and severe CA3 damage) is yet unknown. Further, hippocampal sclerosis is not a pre-requisite for drug-resistant MTLE: studies report an incidence rage of 56-70% in medically intractable patients referred to surgical hippocampal ablation [13]. Thus, while current studies suggest that a fair portion of drug-resistant MTLE patients might benefit of the proposed approach, its feasibility and potential efficacy should be informed by the functional evaluation of the CA3 via intracranial EEG. In the case of severe compromise of the CA3, the mapped statistical parameters of interictal activity (*cf*. **Figure 8)** may help devise the stimulation policy of closed-loop devices relying on seizure prediction and machine learning algorithms fed with those parameters as a vast set of choices for adaptive DBS.

## Ethics statement

This study was conducted in accordance with the National Legislation (D.Lgs. 26/2014) and the European Directive 2010/63/EU and approved by the Institutional Ethics Committee of Istituto Italiano di Tecnologia and by the Italian Ministry of Health (protocol code 860/215-PR, approval date 24/08/2015; protocol code 176AA.NTN9, approval date 20/10/2018).

## CRediT AUTHORSHIP CONTRIBUTION STATEMENT

Davide Caron: formal analysis, investigation, validation. **Stefano Buccelli**: investigation, methodology, software, validation, writing – review & editing. **Ángel Canal-Alonso**: formal analysis, methodology, software, validation, writing – review & editing. **Javad Farsani**: investigation, software (DSP-based closed-loop stimulation bridge-50%), validation. Giacomo Pruzzo: conceptualization – hardware, writing – review & editing. **Bernabé Linares Barranco**: funding acquisition, investigation, software (DSP-based closed-loop stimulation bridge-50%), project administration, resources, supervision, validation, writing – review & editing, **Juan Manuel Corchado**: formal analysis, funding acquisition project administration, resources, supervision, validation, writing – review & editing. **Michela Chiappalone**: conceptualization, funding acquisition, methodology, project administration, resources, supervision, validation, writing – review & editing. **Gabriella** Panuccio: conceptualization, data curation, formal analysis, funding acquisition, investigation, methodology, project administration, resources, software, supervision, validation, visualization, writing – original draft, review & editing.

## Acknowledgements

We thank the Animal Facility technical staff for their help; Alessandra Sanna and Cinzia Nasso for administrative assistance.

## References

[1] Nagaraj V, Lee ST, Krook-Magnuson E, Soltesz I, Benquet P, Irazoqui PP, et al. Future of Seizure Prediction and Intervention: Closing the Loop. Journal of Clinical Neurophysiology. 2015;32(3).

[2] Wagenaar DA, Madhavan R, Pine J, Potter SM. Controlling bursting in cortical cultures with closed-loop multi-electrode stimulation. J Neurosci. 2005;25(3):680–8.

[3] Ben-Menachem E, Krauss GL. Epilepsy: responsive neurostimulation-modulating the epileptic brain. Nature reviews Neurology. 2014;10(5):247–8.

[4] Morrell MJ, Halpern C. Responsive Direct Brain Stimulation for Epilepsy. Neurosurgery clinics of North America. 2016;27(1):111–21.

[5] Pineau J, Guez A, Vincent R, Panuccio G, Avoli M. Treating epilepsy via adaptive neurostimulation: a reinforcement learning approach. International journal of neural systems. 2009;19(4):227–40.

[6] Panuccio G, Guez A, Vincent R, Avoli M, Pineau J. Adaptive control of epileptiform excitability in an in vitro model of limbic seizures. Experimental neurology. 2013;241:179–83.

[7] Kuhlmann L, Lehnertz K, Richardson MP, Schelter B, Zaveri HP. Seizure prediction — ready for a new era. Nature Reviews Neurology. 2018;14(10):618–30.

[8] Sandler RA, Song D, Hampson RE, Deadwyler SA, Berger TW, Marmarelis VZ. Hippocampal closed-loop modeling and implications for seizure stimulation design. Journal of Neural Engineering. 2015;12(5):056017.

[9] Geller EB, Skarpaas TL, Gross RE, Goodman RR, Barkley GL, Bazil CW, et al. Brain-responsive neurostimulation in patients with medically intractable mesial temporal lobe epilepsy. Epilepsia. 2017;58(6):994–1004.

[10] Guggenmos DJ, Azin M, Barbay S, Mahnken JD, Dunham C, Mohseni P, et al. Restoration of function after brain damage using a neural prosthesis. Proc Natl Acad Sci U S A. 2013;110(110):21177–82.

[11] Collinger JL, Foldes S, Bruns TM, Wodlinger B, Gaunt R, Weber DJ. Neuroprosthetic technology for individuals with spinal cord injury. J Spinal Cord Med. 2013;36(36):258–72.

[12] Blümcke I, Thom M, Aronica E, Armstrong DD, Bartolomei F, Bernasconi A, et al. International consensus classification of hippocampal sclerosis in temporal lobe epilepsy: A Task Force report from the ILAE Commission on Diagnostic Methods. Epilepsia. 2013;54(7):1315–29.

[13] Prayson RA. 25 - Pathology of Epilepsy. In: Perry A, Brat DJ, editors. Practical Surgical Neuropathology: A Diagnostic Approach (Second Edition): Elsevier; 2018. p. 617–32.

[14] Benini R, Avoli M. Rat subicular networks gate hippocampal output activity in an in vitro model of limbic seizures. J Physiol. 2005;566(Pt 3):885–900.

[15] Fisher PD, Sperber EF, Moshe SL. Hippocampal sclerosis revisited. Brain & development. 1998;20(20):563–73.

[16] Wozny C, Kivi A, Lehmann T-N, Dehnicke C, Heinemann U, Behr J. Comment on “On the Origin of Interictal Activity in Human Temporal Lobe Epilepsy in Vitro”. Science. 2003;301(301):463.

[17] Knopp A, Kivi A, Wozny C, Heinemann U, Behr J. Cellular and network properties of the subiculum in the pilocarpine model of temporal lobe epilepsy. Journal of Comparative Neurology. 2005;483(483):476–88.

[18] Barbarosie M, Avoli M. CA3-driven hippocampal-entorhinal loop controls rather than sustains in vitro limbic seizures. J Neurosci. 1997;17(17):9308–14.

[19] D’Arcangelo G, Panuccio G, Tancredi V, Avoli M. Repetitive low-frequency stimulation reduces epileptiform synchronization in limbic neuronal networks. Neurobiol Dis. 2005;19(1-2):119–28.

[20] Caron D, Canal-Alonso A, Panuccio G. Mimicking CA3 Temporal Dynamics Controls Limbic Ictogenesis. Biology. 2022;11(3).

[21] Avoli M, D’Antuono M, Louvel J, Kohling R, Biagini G, Pumain R, et al. Network and pharmacological mechanisms leading to epileptiform synchronization in the limbic system in vitro. Progress in neurobiology. 2002;68(68):167–207.

[22] McIntyre DC, Gilby KL. Mapping seizure pathways in the temporal lobe. Epilepsia. 2008;49 Suppl 3:23–30.

[23] Bartolomei F, Khalil M, Wendling F, Sontheimer A, Regis J, Ranjeva JP, et al. Entorhinal cortex involvement in human mesial temporal lobe epilepsy: an electrophysiologic and volumetric study. Epilepsia. 2005;46(46):677–87.

[24] Spencer SS, Spencer DD. Entorhinal-hippocampal interactions in medial temporal lobe epilepsy. Epilepsia. 1994;35(35):721–7.

[25] Bartolomei F, Chauvel P, Wendling F. Epileptogenicity of brain structures in human temporal lobe epilepsy: a quantified study from intracerebral EEG. Brain. 2008;131(Pt 7):1818–30.

[26] Panuccio G, Colombi I, Chiappalone M. Recording and Modulation of Epileptiform Activity in Rodent Brain Slices Coupled to Microelectrode Arrays. J Vis Exp. 2018(135).

[27] Rutecki PA, Lebeda FJ, Johnston D. 4-Aminopyridine produces epileptiform activity in hippocampus and enhances synaptic excitation and inhibition. J Neurophysiol. 1987;57(57):1911–24.

[28] Panuccio G, D’Antuono M, de Guzman P, De Lannoy L, Biagini G, Avoli M. In vitro ictogenesis and parahippocampal networks in a rodent model of temporal lobe epilepsy. Neurobiol Dis. 2010;39(39):372–80.

[29] Daubechies I. Ten lectures on wavelets: Society for Industrial and Applied Mathematics; 1992. 357 p.

[30] Toprani S, Durand DM. Fiber tract stimulation can reduce epileptiform activity in an in-vitro bilateral hippocampal slice preparation. Experimental neurology. 2013;240:28–43.

[31] Mizuseki K, Buzsaki G. Preconfigured, skewed distribution of firing rates in the hippocampus and entorhinal cortex. Cell reports. 2013;4(4):1010–21.

[32] Buzsaki G, Mizuseki K. The log-dynamic brain: how skewed distributions affect network operations. Nature reviews Neuroscience. 2014;15(15):264–78.

[33] Hintze JL, Nelson RD. Violin Plots: A Box Plot-Density Trace Synergism. The American Statistician. 1998;52(52):181–4.

[34] Seth AK, Barrett AB, Barnett L. Granger causality analysis in neuroscience and neuroimaging. J Neurosci. 2015;35(35):3293–7.

[35] Shils JL, Alterman RL, Arle JE. Deep Brain Stimulation Fault Testing. In: Tarsy D, Vitek JL, Starr PA, Okun MS, editors. Deep Brain Stimulation in Neurological and Psychiatric Disorders. Totowa, NJ: Humana Press; 2008. p. 473–94.

[36] Sohal VS, Sun FT. Responsive Neurostimulation Suppresses Synchronized Cortical Rhythms in Patients with Epilepsy. Neurosurgery clinics of North America. 2011;22(22):481–8.

[37] Ronborg SN, Esteller R, Tcheng TK, Greene DA, Morrell MJ, Wesenberg Kjaer T, et al. Acute effects of brain-responsive neurostimulation in drug-resistant partial onset epilepsy. Clin Neurophysiol. 2021;132(132):1209–20.

[38] Bush K, Panuccio G, Avoli M, Pineau J. Evidence-based modeling of network discharge dynamics during periodic pacing to control epileptiform activity. J Neurosci Methods. 2012;204(204):318–25.

[39] Paschen E, Elgueta C, Heining K, Vieira DM, Kleis P, Orcinha C, et al. Hippocampal low-frequency stimulation prevents seizure generation in a mouse model of mesial temporal lobe epilepsy. Elife. 2020;9.

[40] Colom LV, Saggau P. Spontaneous interictal-like activity originates in multiple areas of the CA2-CA3 region of hippocampal slices. J Neurophysiol. 1994;71(71):1574–85.

[41] Wittner L, Miles R. Factors defining a pacemaker region for synchrony in the hippocampus. J Physiol. 2007;584(Pt 3):867–83.

[42] Barbarosie M, Louvel J, Kurcewicz I, Avoli M. CA3-released entorhinal seizures disclose dentate gyrus epileptogenicity and unmask a temporoammonic pathway. J Neurophysiol. 2000;83(83):1115–24.

[43] Swartzwelder HS, Lewis DV, Anderson WW, Wilson WA. Seizure-like events in brain slices: suppression by interictal activity. Brain Res. 1987;410(410):362–6.

[44] Bragdon AC, Kojima H, Wilson WA. Suppression of interictal bursting in hippocampus unleashes seizures in entorhinal cortex: a proepileptic effect of lowering [K+]o and raising [Ca2+]o. Brain Res. 1992;590(1-2):128–35.

[45] Buonomano DV, Maass W. State-dependent computations: spatiotemporal processing in cortical networks. Nature reviews Neuroscience. 2009;10(10):113–25.

[46] Chvojka J, Kudlacek J, Chang WC, Novak O, Tomaska F, Otahal J, et al. The role of interictal discharges in ictogenesis - A dynamical perspective. Epilepsy Behav. 2021;121(Pt B):106591.

[47] Dorn T, Witte OW. Refractory periods following interictal spikes in acute experimentally induced epileptic foci. Electroencephalogr Clin Neurophysiol. 1995;94(94):80–5.

[48] Dzhala VI, Staley KJ. Transition from Interictal to Ictal Activity in Limbic Networks In Vitro. The Journal of Neuroscience. 2003;23(23):7873–80.

[49] Staley KJ, Dudek FE. Interictal spikes and epileptogenesis. Epilepsy Curr. 2006;6(6):199–202.

[50] Chang WC, Kudlacek J, Hlinka J, Chvojka J, Hadrava M, Kumpost V, et al. Loss of neuronal network resilience precedes seizures and determines the ictogenic nature of interictal synaptic perturbations. Nat Neurosci. 2018;21(21):1742–52.

[51] Lai N, Cheng H, Li Z, Wang X, Ruan Y, Qi Y, et al. Interictal-period-activated neuronal ensemble in piriform cortex retards further seizure development. Cell reports. 2022;41(41):111798.

[52] Lai N, Li Z, Xu C, Wang Y, Chen Z. Diverse nature of interictal oscillations: EEG-based biomarkers in epilepsy. Neurobiol Dis. 2023;177:105999.

[53] Celiker Uslu S, Yuksel B, Tekin B, Sariahmetoglu H, Atakli D. Cognitive impairment and drug responsiveness in mesial temporal lobe epilepsy. Epilepsy Behav. 2019;90:162–7.

[54] Blumcke I, Thom M, Aronica E, Armstrong DD, Bartolomei F, Bernasconi A, et al. International consensus classification of hippocampal sclerosis in temporal lobe epilepsy: a Task Force report from the ILAE Commission on Diagnostic Methods. Epilepsia. 2013;54(54):1315–29.

