## Supplementary figures and table for "Biohybrid restoration of the hippocampal loop re-establishes the non-seizing state in an in vitro model of limbic seizures"

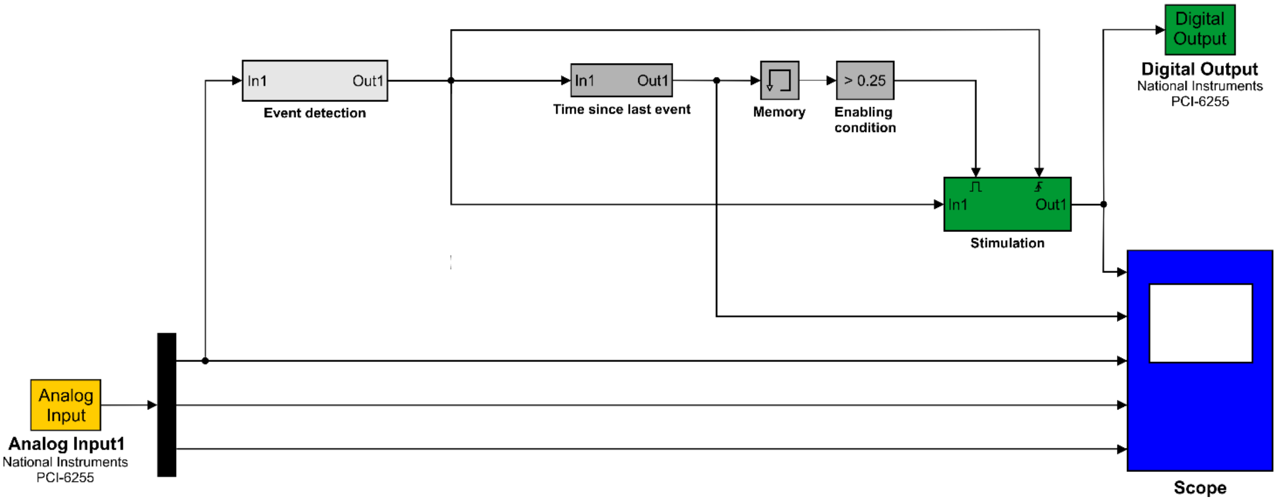


*Supplementary Figure 1 – Simulink model for biohybrid bridging.*

The model consists of input (yellow), processing (grey) and output (green) blocks, along with a scope window (blue). The processing blocks include routines for the event detection (light grey) and the decision to stimulate (dark grey). The Analog Input block retrieves the signals from the MEA system and feeds the signal of one selected electrode from the CA3 to the Event detection block. The latter implements a hard-threshold crossing algorithm and feeds its output to the Time since last event block to start the decision routine. The Stimulation block is triggered if the enabling condition is met. The stimulation block then triggers the Digital Output block, which, in turn, activates the TTL signal to the external stimulator. The Scope window enables the real-time visualization of the MEA signals, as well as the output of the *Event detection*, *Time since last event*, and *Stimulation* blocks.

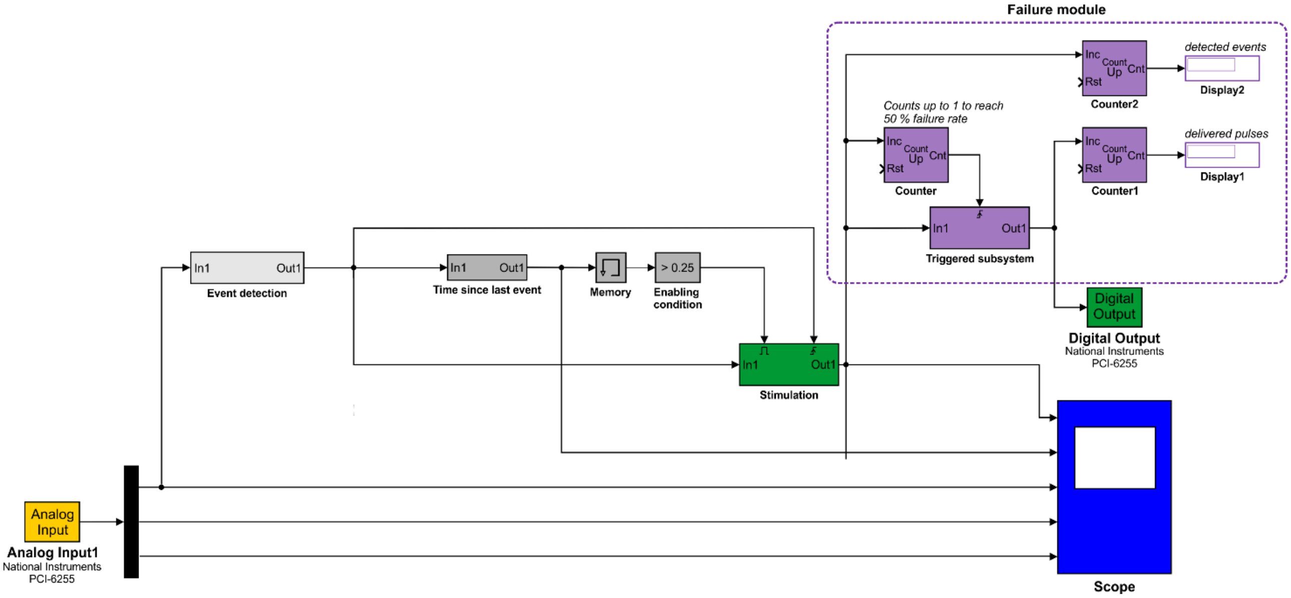


*Supplementary Figure 2 – Simulink model implementation for 50% failure testing.*

The model is based on the same architecture depicted in **Supplementary Figure 1** with the addition of a failure module (purple blocks, framed by the purple dashed box), based on a counter of triggered events.

*Supplementary Table 1 – Bridge performance overview, assessed with Granger causality (data from Figure 10B).*

| CA3-to-CTX | Disconnected | Bridge-bw5 | Bridge-50% | Bridge-100% | Connected |
|---|---|---|---|---|---|
| Disconnected | | n.s. | *** | *** | *** |
| Bridge-bw5 | n.s. | | ** | ** | *** |
| Bridge-50% | ** | ** | | * | *** |
| Bridge-100% | ** | ** | * | | n.s. |
| Connected | *** | *** | *** | n.s. | |

| CTX-to-CA3 | Disconnected | Bridge-bw5 | Bridge-50% | Bridge-100% | Connected |
|---|---|---|---|---|---|
| Disconnected | | n.s. | *** | *** | *** |
| Bridge-bw5 | n.s. | | ** | ** | *** |
| Bridge-50% | *** | ** | | * | ** |
| Bridge-100% | *** | ** | * | | n.s. |
| Connected | *** | *** | ** | n.s. | |

* $p < 0.05$; ** $p < 0.01$; *** $p < 0.001$; n.s.: not significant.
